# Correlation of Computerized Tomography (CT) Severity Score for COVID-19 pneumonia with Clinical Outcomes

**DOI:** 10.1101/2021.01.15.426787

**Authors:** Kiran Hilal, Jehanzeb Shahid, Abdullah Ameen, Russell Martins, Avinash Nankani, Ainan Arshad

## Abstract

**Introduction:** Various CT severity scores have already been described in literature since the start of this pandemic. One pertinent issue with all of the previously described severity scores is their relative challenging calculation and variance in inter-observer agreement. The severity score proposed in our study is relatively simpler, easier to calculate and apart from a trained radiologist, can easily be calculated even by physicians with good inter-observer agreement. Therefore, a rapid CT severity score calculation can give a clue to physician about possible clinical outcome without being dependent on radiologist who may not be readily available especially in third world countries.

**Objective:** The objective of this study is to develop a simple CT severity score (CT-SS) with good inter-observer agreement and access its correlation with clinical outcome.

**Methods:** This retrospective study was conducted by the Department of Radiology and Internal Medicine, at the Aga Khan University Hospital Karachi, from April 2020 to August 2020. Non-probability consecutive sampling was used to include all patients who were positive for COVID-19 on PCR, and underwent CT chest examination at AKUH. Severity of disease was calculated in each lobe on the basis of following proposed CT severity scoring system (CT-SS). For each lobe the percentage of involvement by disease was scored – 0% involvement was scored 0, <50% involvement was scored 1 and >50% involvement was scored 2. Maximum score for one lobe was 2 and hence total maximum overall score for all lobes was 10. Continuous data was represented using mean and standard deviation, and compared using independent sample t-tests. Categorical data was represented using frequencies and percentages, and compared using Chi-squared tests. Inter-observer reliability between radiologist and COVID intensivist for the 10 point CT-SS rated on 0-10 was assessed using the Kappa statistic. A p-value < 0.05 was considered significant for all analyses.

**Results:** A total of 73 patients were included, the majority male (58.9%) with mean age 55.8 ± 13.93 years. The CT-SS rated on 0-10 showed substantial inter-observer reliability between radiologist and intensivist with a Kappa statistic of 0.78. Patients with CT-SS 8-10 had a significantly higher ICU admission & intubation rate (53.8% vs. 23.5%) and mortality rate (35.9% vs. 11.8%; p = 0.017), as compared to those with CT-SS 0-7.

**Conclusion:** We conclude that the described CT severity score (CT-SS) is a quick, effective and easily reproducible tool for prediction of adverse clinical outcome in patients with COVID 19 pneumonia. The tool shows good inter-observer agreement when calculated by radiologist and physician independently.

## Introduction

The World Health Organization (WHO) officially declared COVID 19 (coronavirus disease 2019) as a pandemic on March 11, 2020 and required international action in four key areas: to prepare and be ready; detect, protect and treat; reduce transmission; and innovate and learn ^(1)^. At the time of writing this article, there are more than 25 million confirmed cases of COVID-19 with more than 800000 death worldwide ^(2)^.

COVID-19 is highly transmissible but with a relatively low mortality rate with (1% − 3.5%), except in older individuals with multiple underlying health conditions ^(3)^. Most patients present with symptoms of fever, cough, dyspnea and fatigue, with only 15% − 20% developing severe pneumonia and 5% − 10% requiring critical care ^(3)^. Treatment of COVID 19 varies as per the severity of symptoms, with severe cases needing intensive care admission. While reverse transcriptase polymerase chain reaction (RT-PCR) is the most reliable means for diagnosis of COVID 19 disease ^(4)^, computed tomography (CT) also has a high sensitivity for the diagnosis of COVID-19. Most common CT findings in patient with COVID 19 are peripherally distributed multi focal ground glass haziness and consolidations ^(4-6)^.

In the early phase of COVID-19, clinical and imaging features are most important in establishment of the diagnosis, evaluation of changes in severity, and adjustment of treatment plan. CT scan chest has a very high sensitivity for detection of COVID 19 pneumonia and has been reported up to 99% in Wuhan, China and worldwide has been reported from 61% to 98% ^(7)^. The reported sensitivity of chest x ray for detection of COVID 19 pneumonia is 73% for the initial radiograph and increasing further with time and has sensitivity of up to 83% for the follow-up radiographs ^(8)^.

Various CT severity scores have already been described in literature since the start of this pandemic ^(15-17)^. One pertinent issue with all of the previously described severity scores is their relative challenging calculation and variance in inter-observer agreement. The severity score proposed in our study is relatively simpler, easier to calculate and apart from a trained radiologist, can easily be calculated even by physicians with good inter-observer agreement. Therefore, a rapid CT severity score calculation can give a clue to physician about possible clinical outcome without being dependent on radiologist who may not be readily available especially in third world countries. The objective of this study is to develop a simple CT severity score (CT-SS) with good inter-observer agreement and access its correlation with clinical outcome.

## Methods

### Setting and Sample

This retrospective study was conducted by the Department of Radiology and Internal Medicine, at the Aga Khan University Hospital (AKUH), Karachi, from April 2020 to August 2020. Ethical approval was received from the institutional review board. Non-probability consecutive sampling was used to include all patients who were positive for COVID-19 on PCR, and underwent CT chest examination at AKUH. Patients with cardiac failure and fluid overload, metastatic lung disease, non COVID pulmonary pneumonia, recent history of cardiothoracic surgery and patients with history of road traffic accident with pulmonary hemorrhage were excluded from the study.

### Proposed CT severity scoring system

Severity of disease was calculated in each lobe on the basis of following proposed CT severity scoring system (Fig. 1,2,3).

**Fig 1.**
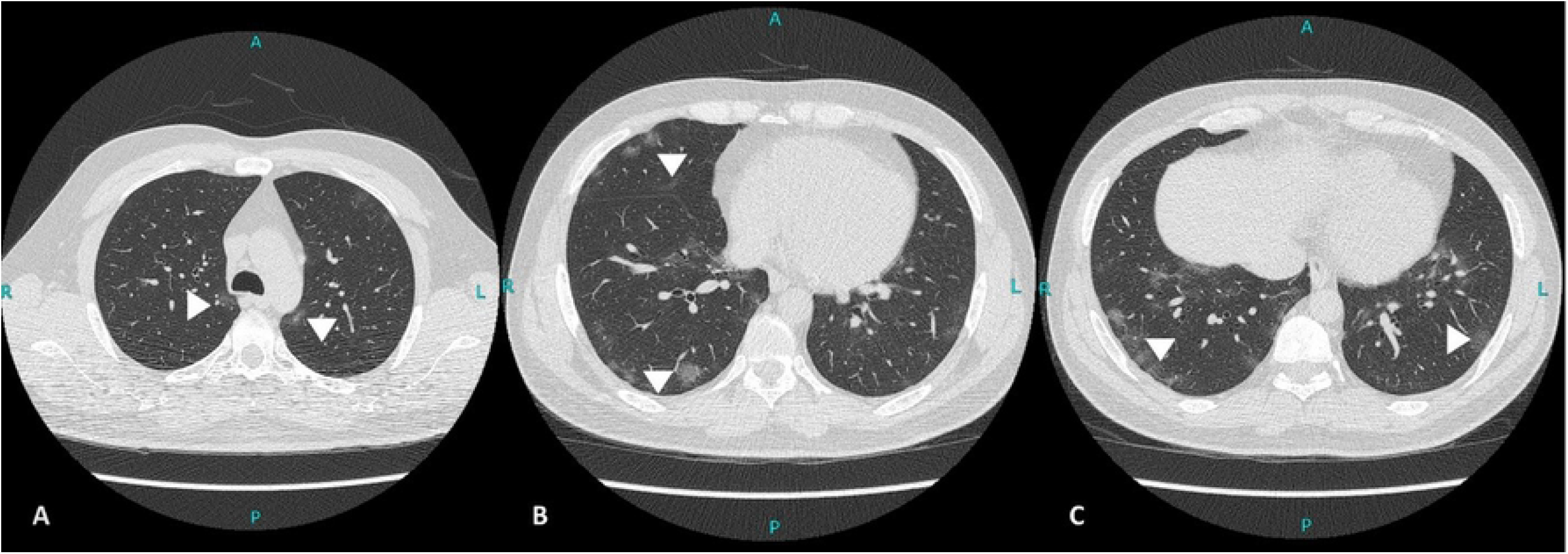
(A-C): Unenhanced axial sections from lung window of CT chest of 42 years old male patient with PCR confirmed COVID-19 pneumonia. Multiple scattered areas of mostly peripheral ground glass opacities in both lungs involving bilateral upper, right mid and bilateral lower lung lobes (white arrows). All lobes are showing <50% parenchymal involvement. CT severity score (CTSS) calculated as 5.

**Fig 2.**
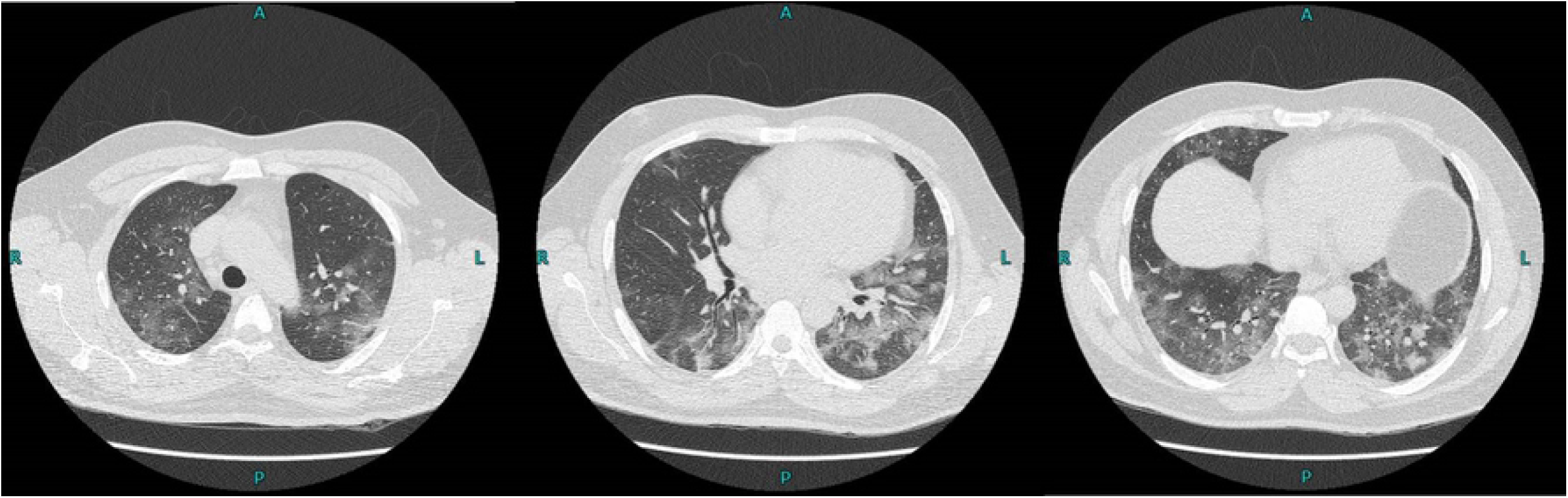
(A-C) Axial sections from CT chest lung window of a 59 years old male patient with PCR confirmed COVID-19 pneumonia. Extensive ground glass opacification in both lung fields showing >50% involvement of all the lung lobes. CT severity score(CTSS) calculated as 10.

**Fig 3.**
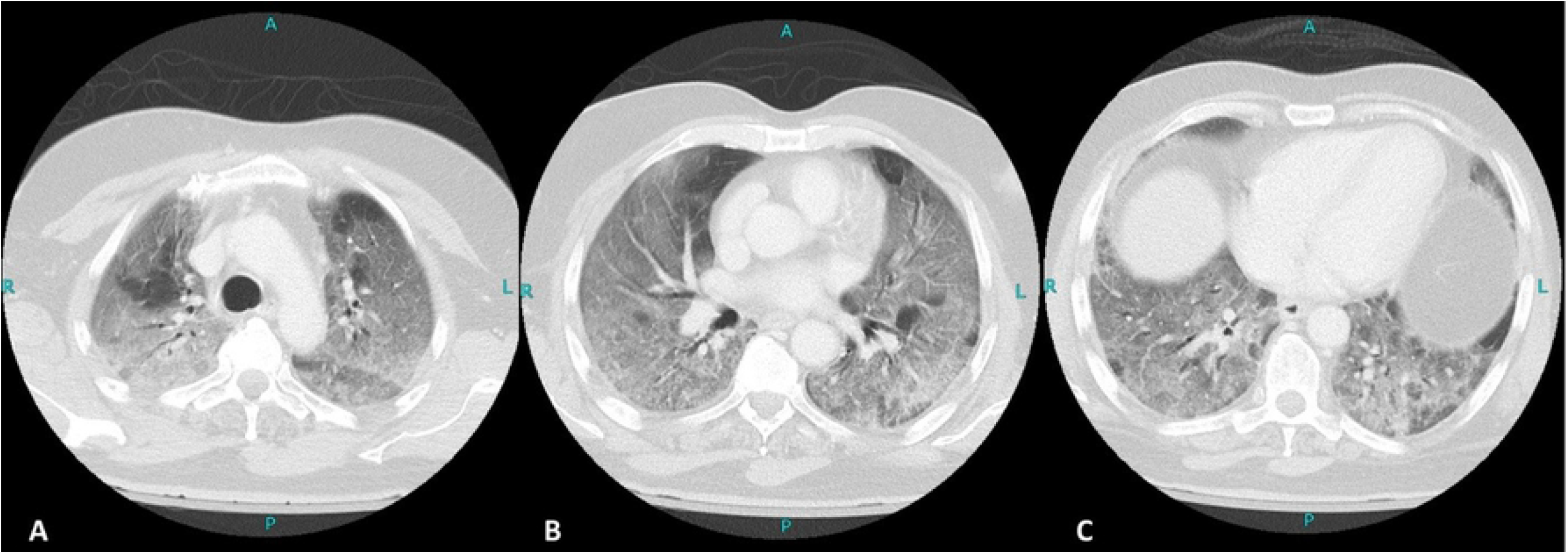
(A-C) Axial sections from CT chest lung window of a 59 years old male patient with PCR confirmed COVID-19 pneumonia. Extensive ground glass opacification in both lung fields showing >50% involvement of all the lung lobes. CT severity score(CTSS) calculated as 10.

**Fig 3.**
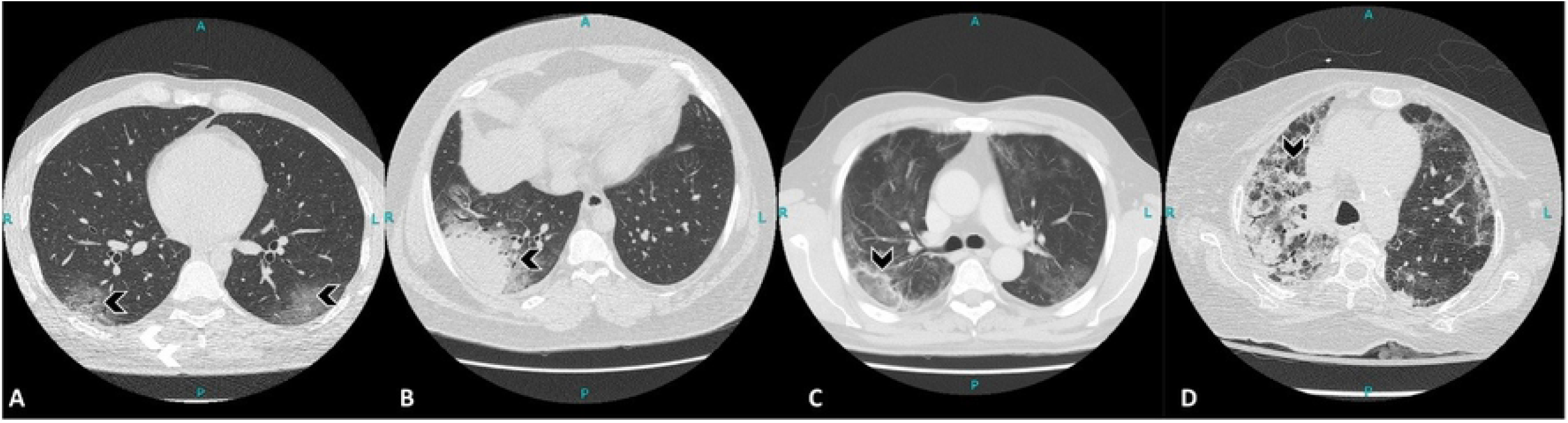
(A-D) :Multiple axial sections of CT chest lung windows of various patients showing imaging spectrum of COVID-19 pneumonia (black arrowheads). Bilateral ground glass opacities (A), consolidation (B), Atoll sign (C), crazy paving pattern (D).

For each lobe the percentage of involvement by disease was scored – 0% involvement was scored 0, <50% involvement was scored 1 and >50% involvement was scored 2. Maximum score for one lobe was 2 and hence total maximum score for right lung was 6 and for left lung total maximum score was 4. Total maximum overall score for all lobes was 10.

### Data Collection

Patients’ data was retrieved from the hospital electronic medical records. CT scan images were reviewed independently by two radiologists from the radiological database. One radiologist had more than eight years and other radiologist had more than five years of post-consultant experience in chest imaging. For the purpose of analysis, CT Severity Score (CT-SS) on 0-10 were categorized as Low (0-7) and High (8-10), by both radiologists independently. For cases with difference in calculated score by two radiologists, a consensus was reached for calculated score in post score calculation meeting. CT-SS was independently calculated by a COVID intensivist with about six months of experience in COVID ICU, who was blinded with CT-SS calculated by radiologists. Inter-observer reliability between radiologist and COVID intensivist for the 10 point CT-SS rated on 0-10 was assessed using the Kappa statistic. All clinical variables extracted included patient demographics, clinical characteristics, laboratory readings, and hospitalization outcomes.

### Data Analysis

Data analysis was performed using IBM SPSS v. 21. Continuous data was represented using mean and standard deviation, and compared using independent sample t-tests. Categorical data was represented using frequencies and percentages, and compared using Chi-squared tests. A p-value < 0.05 was considered significant for all analyses.

## Results

A total of 73 patients were included, the majority male (58.9%) with mean age 55.8 ± 13.93 years. The most common comorbidities were hypertension (38.4%) and type 2 diabetes mellitus (28.8%), while the most common presenting complaints were shortness of breath (53.4%) and cough (41.1%). 39.7% patients required ICU admission and intubation, and 56.2% had a length of hospital stay greater than 10 days. The mortality rate in this study was 24.7%.

The CT-SS rated on 0-10 showed substantial inter-observer reliability between radiologist and COVID intensivist with a Kappa statistic of 0.78. When categorized according to CT-SS, around half fell into the High (8-10) category (53.4%), with the rest into the Low (0-7) category. Patients with a High CT-SS were more likely to present with shortness of breath (71.8% vs. 32.4%; p = 0.004), cough (56.4% vs. 23.5%; p = 0.001) and fever (59% vs. 32.4%; p = 0.023). Moreover, patients in the 8-10 category had a higher mean C-reactive protein (109.14 ± 86.89 vs. 62.80 ± 67.59; p = 0.022) and lactate dehydrogenase (515.27 ± 238.53 vs. 380.50 ± 149.09; p = 0.036), than patients in the 0-7 category. Patients with CT-SS 8-10 had a significantly higher ICU admission & intubation rate (53.8% vs. 23.5%) and mortality rate (35.9% vs. 11.8%; p = 0.017), as compared to those with CT-SS 0-7.

**Flow Chart 1.**
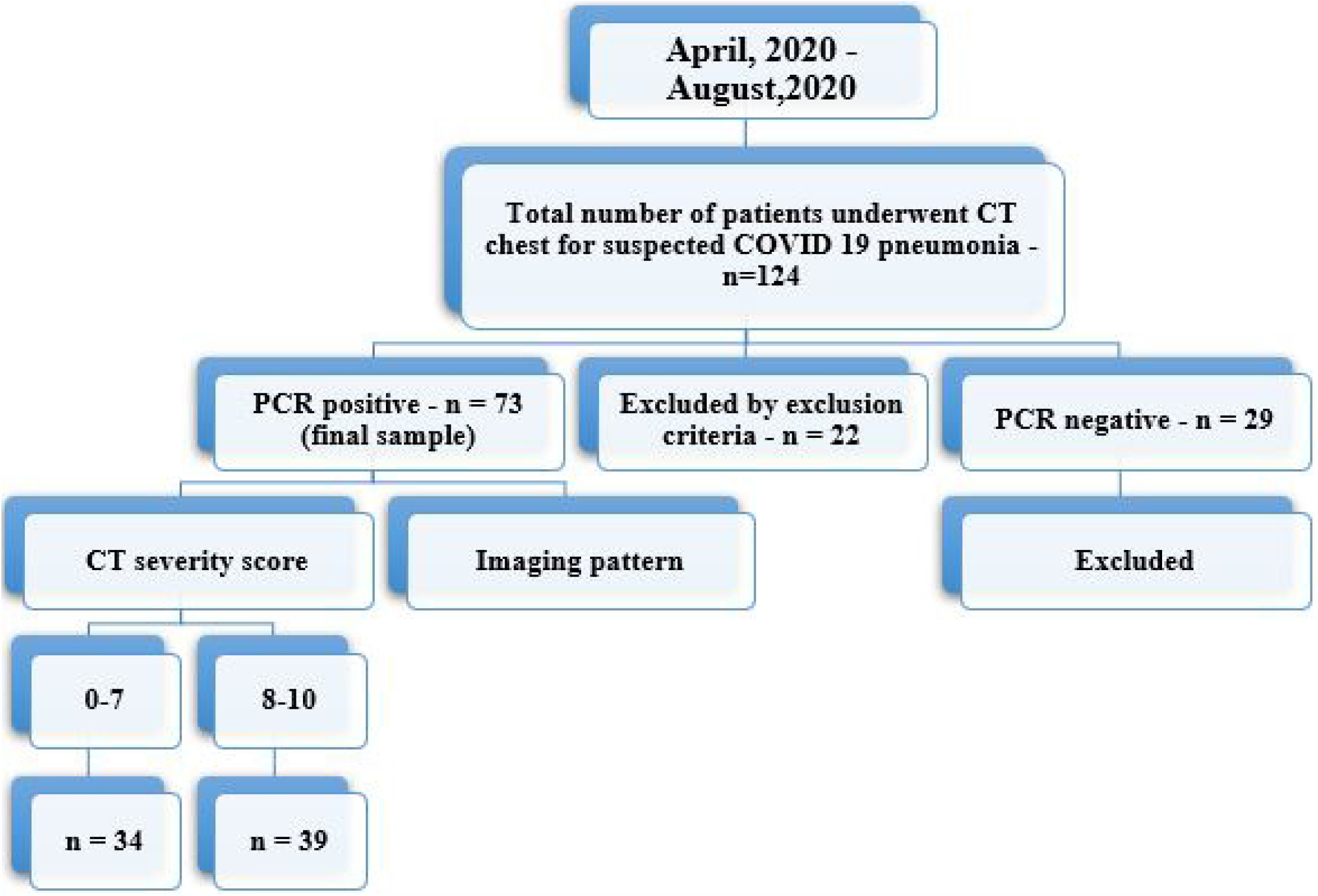
Patient selection strategy and final sample size (n = number of patients)

The distribution of CT scan findings is shown in **Table 2**. Most common imaging features included ground glass opacities, consolidations, crazy paving and atoll sign(Fig. 3). Patients with CT-SS 8-10 were significantly more likely to have ground glass opacities, interstitial infiltrates, crazy paving pattern, sub-pleural disease, anterior disease, posterior disease, and involvement of all lung lobes (all p < 0.05).

**Table 1:** Clinical Characteristics of Patients. COPD: Chronic Obstructive Pulmonary Disease; ICU: Intensive Care Unit; LOS: Length of Stay

| Variable | Overall (N=73)<br>n (%) / Mean ± SD | CT-SS | | P-Value |
| --- | --- | --- | --- | --- |
|  |  | 0-7 (N=34)<br>n (%) | 8-10 (N=39)<br>n (%) |  |
| Age (years) | 55.8 ± 13.93 | 53.82 ± 16.34 | 57.54 ± 11.37 | 0.259 |
| Gender |  |  |  |  |
| Male | 43 (58.9) | 20 (58.8) | 23 (59.0) | 0.990 |
| Female | 30 (41.1) | 14 (41.2) | 16 (41.0) |  |
| Comorbid |  |  |  |  |
| Hypertension | 28 (38.4) | 16 (47.1) | 12 (30.8) | 0.153 |
| Type 2 Diabetes Mellitus | 21 (28.8) | 8 (23.5) | 13 (33.3) | 0.356 |
| Ischemic Heart Disease | 10 (13.7) | 6 (17.6) | 4 (10.3) | 0.360 |
| Chronic Kidney Disease | 3 (4.1) | 2 (5.9) | 1 (2.6) | 0.476 |
| COPD/Asthma | 1 (1.4) | 1 (2.9) | 0 (0) | > 0.466 |
| C-Reactive Protein | 87.46 ± 81.25 | 62.80 ± 67.59 | 109.14 ± 86.89 | **0.022** |
| Procalcitonin (ng/mL) | 0.55 ± 0.89 | 0.59 ± 0.84 | 0.51 ± 0.96 | 0.736 |
| Lactate Dehydrogenase | 451.09 ± 209.96 | 380.50 ± 149.09 | 515.27 ± 238.53 | **0.036** |
| D-Dimer | 4.77 ± 4.96 | 4.07 ± 3.55 | 5.32 ± 5.85 | 0.440 |
| Symptoms |  |  |  |  |
| Shortness of Breath | 39 (53.4) | 11 (32.4) | 28 (71.8) | **0.004** |
| Cough | 30 (41.1) | 8 (23.5) | 22 (56.4) | **0.001** |
| Fever | 34 (46.6) | 11 (32.4) | 23 (59.0) | **0.023** |
| Chest Pain/Discomfort | 5 (6.8) | 3 (8.8) | 2 (5.1) | 0.533 |
| No symptoms | 16 (21.9) | 11 (32.4) | 5 (12.8) | 0.044 |
| LOS>10 Days | 41 (56.2) | 15 (44.1) | 26 (66.7) | 0.053 |
| ICU & Intubation | 29 (39.7) | 8 (23.5) | 21 (53.8) | **0.008** |
| Outcome |  |  |  |  |
| Recovered | 55 (75.3) | 30 (88.2) | 25 (64.1) | **0.017** |
| Mortality | 18 (24.7) | 4 (11.8) | 14 (35.9) |  |

**Table 2:** CT Characteristics

| Table 2: CT Characteristics |  |  |  |  |
| --- | --- | --- | --- | --- |
| Radiological Properties | Overall (N=73)<br>n (%) | CT-SS |  | P-Value |
|  |  | 0-7 (N=34)<br>n (%) | 8-10 (N=39)<br>n (%) |  |
| GGO | 66 (90.4) | 28 (82.4) | 38 (97.4) | **0.045** |
| Consolidation | 51 (69.9) | 20 (58.8) | 31 (79.5) | 0.055 |
| Interstitial Infiltrates | 32 (43.8) | 7 (20.6) | 25 (64.1) | **< 0.001** |
| Crazy Paving Pattern | 22 (30.1) | 3 (8.8) | 19 (48.7) | **< 0.001** |
| Atoll Sign | 7 (9.6) | 2 (5.9) | 5 (12.8) | 0.438 |
| Subpleural Disease | 68 (93.2) | 20 (85.3) | 39 (100) | **0.019** |
| Perihilar Disease | 42 (57.5) | 9 (26.5) | 33 (84.6) | **< 0.001** |
| Anterior Disease | 60 (82.2) | 21 (61.8) | 39 (100) | **< 0.001** |
| Posterior Disease | 69 (94.5) | 30 (88.2) | 39 (100) | **0.043** |
| Right Upper Lobe Involved | 64 (87.7) | 25 (73.5) | 39 (100) | **0.001** |
| Right Middle Lobe Involved | 61 (83.6) | 22 (64.7) | 39 (100) | **< 0.001** |
| Right Lower Lobe Involved | 65 (89.0) | 26 (76.5) | 39 (100) | **0.001** |
| Left Upper Lobe Involved | 66 (90.4) | 27 (79.4) | 39 (100) | **0.003** |
| Left Lower Lobe Involved | 66 (90.4) | 28 (82.4) | 28 (97.4) | **0.045** |

## Discussion

In our hospital, when patients with clinical symptoms suggestive of COVID-19 infection (such as shortness of breath, cough, and fever) are received, they are initially resuscitated and worked up with laboratory tests and chest radiograph. If radiographic findings reveal suspicious infiltrates, or there is any history of contact with a COVID-19 individual, a nasopharyngeal swab for RT-PCR is performed. However, the results of the RT-PCR arrive after 8-12 hours or even longer if sent at night or on a weekend, sacrificing precious time in the management of a patient.

COVID 19 pneumonia has so far shown very unpredictable clinical course with clinical features ranging from asymptomatic to severe disease with ARDS and multi-organ dysfunction. Currently there is no approved biomarker which can predict the disease course, identify the group of patients who need urgent medical care, hospitalization and estimate the associated mortality ^(9)^.

We started this study with a hypothesis of correlating the CT, laboratory and clinical features of the COVID 19 pneumonia and predict the disease outcome and to categorize the disease as mild or severe depending on the CT features by developing a CT severity scoring system which is easy to use, reproducible and can also be used by the primary treating medical specialist.

Ground glass opacification with or without consolidations is the hallmark feature of COVID 19 pneumonia as reported by the prior studies ^(10, 11)^ and is also confirmed by our present study. As reported previously, the late or severe stage of disease is led by activation of humoral and cellular immunity mediated by virus specific B and T cells. The higher prevalence of consolidation and crazy paving pattern in our study is explained by the above mentioned mechanism of injury which likely is due to commutation of pulmonary edema, superadded bacterial infection and interstitial inflammatory changes ^(12)^.

The main goal which we achieved with this study was to develop and validate an easy and reproducible CT severity scoring system by combining the clinical, laboratory and CT features of COVID 19 pneumonia which is applicable to the population in our region. Our study revealed that the patients with score of 7 or less had shorter length of stay in hospital (<10 days in 44.1% of the patients), lower ICU admissions and need for mechanical ventilation (23.5%), higher recovery (88.2%) and lower mortality (11.8%) rates. However patients with CT severity scores of >7 had longer hospital stay (>10 days in 66.7% of the patients), higher ICU admission and need for mechanical ventilation (53.8%), lower recovery (64.1%) and relatively higher mortality (35.9%) rates compared to the group of patients with CT severity score of 7 or less.

In addition to the CT features, the serum CRP levels were found to be significantly raised in patients with CT severity of >7 (mean value of 87.4 +/− 67.5) as compared to those with CT severity scores of 7 or less (mean value of 109.1 +/− 86.8) with a p-value of 0.022. Serum LDH levels were also found to be significantly raised in patients with severe disease (mean value of 512.2 +/− 238.5) as compared to those with mild disease (mean value of 380.5 +/− 149) with a p-value of 0.036. The serum D-Dimer levels were also increased in severe cases (5.3 +/− 5.8) versus in mild cases (4 +/− 3.5), however not statically signification with p-value of 0.44. These findings have been reported previously and also are confirmed by our study ^(13, 14)^

Various CT severity scores have already been described in literature since the start of this pandemic ^(15-17)^. But one pertinent problem in all of the previously described scores is relatively challenging calculation and variance in inter-observer agreement. The severity score proposed in our study is relatively simpler, easier to calculate and apart from a trained radiologist, can easily be calculated even by physicians with good inter-observer agreement. Our study demonstrated a good interobserver agreement of calculated CT severity score between radiologist and physician with a kappa value of 0.78. The scans with disagreement in severity score were mostly those scans with very subtle ground glass opacities or diseases with interstitium predominant pattern which is probably understandable as a physician is not typically trained for doing regular radiological reporting.

A recent study by Francone M et al showed predominant ground glass opacification in early and crazy paving pattern and consolidation in later phases of disease. This study also revealed higher CRP, D-Dimers levels with increased mortality in patients labeled as severe disease according to their CT severity scoring system ^(15)^.

Another study performed on Chinese population with 102 patients with lobe based CT severity scoring system showed increased total CT severity score in patients with severe disease as compared to the mild cases. The CT severity score was mainly correlated with laboratory findings in this study and clinical feature including mortality and outcome were not included in this study ^(16)^.

Study performed by Zhang J et al on 108 patients revealed positive association of CT severity score with CRP, erythrocyte sedimentation rate (ESR), D-Dimer and procalcitonin and negative association with lymphocyte count ^(17)^.

The findings of our present study show that CT scan can help prognosticate patients and identify those at higher risk of severe outcomes, including mortality. Early intervention, consisting of intensive care specialist consultation, ICU admission, early intubation, steroid treatment, may be performed in patients with CT severity score of 7 or more, to help lower mortality rates in such patients.

Limitations of this study include its small sample size, data from only a single center and unavailability of follow-up CT scans of all patients. The follow up CT were available in limited number of patients and were performed mostly in those patients in whom progression of disease was suspected or a second pathology was also being considered apart from COVID pneumonia.

As a second peak of COVID-19 threatens Pakistan like the rest of the world, further validation studies with larger patient populations using CT-based scoring systems are needed to evaluate the effectiveness of CT scans in the prognostication and management of COVID-19 patients.

## Conclusion

We conclude that the described CT severity score (CT-SS) is a quick, effective and easily reproducible tool for prediction of adverse clinical outcome in patients with COVID 19 pneumonia. The tool shows good inter-observer agreement when calculated by radiologist and physician independently.

## Acknowledgement

None

